# A second generation IL-2 receptor-targeted diphtheria fusion toxin exhibits anti-tumor activity and synergy with anti-PD-1 in melanoma

**DOI:** 10.1101/420158

**Authors:** Laurene S. Cheung, Juan Fu, Pankaj Kumar, Amit Kumar, Michael E. Urbanowski, Elizabeth A. Ihms, Sadiya Parveen, C. Korin Bullen, Garrett Patrick, Robert Harrison, John R. Murphy, Drew M. Pardoll, William R. Bishai

## Abstract

Denileukin diftitox (DAB_1-389_-IL-2, Ontak^®^) is a diphtheria toxin-based fusion protein that depletes CD25-positive cells including regulatory T cells (Tregs) and was approved for the treatment of persistent or recurrent cutaneous T cell lymphoma. However, the clinical use of denileukin diftitox was limited by vascular leak toxicity and production issues related to drug aggregation and purity. We found that a single amino acid substitution (V6A) in a motif associated with vascular leak induction yields a fully active, second-generation biologic, s-DAB_1-386_-IL-2(V6A), which elicits 50-fold less HUVEC monolayer permeation and is 3.7-fold less lethal to mice by LD_50_ analysis than s-DAB_1-386_-IL-2 Additionally, to overcome aggregation problems, we developed a novel production method for the fusion toxin using *Corynebacterium diphtheriae* that secretes fully-folded, biologically active, monomeric s-DAB_1-386_-IL-2 into the culture medium. Using the poorly immunogenic mouse B16F10 melanoma model, we initiated treatment 7 days after tumor challenge and observed that, while both s-DAB_1-386_-IL-2(V6A) and s-DAB_1-386_-IL-2 are inhibitors of tumor growth, the capacity to treat with higher doses of s-DAB_1-386_-IL-2(V6A) could provide a superior activity window. In a sequential dual therapy study in tumors that have progressed for 10 days both s-DAB_1-386_-IL-2(V6A) and s-DAB_1-386_-IL-2 given prior to checkpoint inhibition with anti-PD-1 antibodies inhibited tumor growth, while either drug given as monotherapy had less effect. s-DAB_1-386_-IL-2(V6A), a fully monomeric protein with reduced vascular leak, is a second-generation diphtheria toxin-based fusion protein with promise as a cancer immunotherapeutic both alone and in conjunction with PD-1 blockade.

**Significance Statement:** Regulatory T cells (Tregs) infiltrate tumors in various cancers and promote an immunosuppressive microenvironment that hinders anti-tumor immunity. Denileukin diftitox, a diphtheria toxin-based fusion protein that depletes Tregs, was approved for the treatment of T cell malignancies, but its clinical use was limited due to the presence of protein aggregates and toxicity associated with vascular leakage. Here we report the production of a second generation IL-2 receptor-targeted, fully-folded, monomeric diphtheria fusion toxin, and a V6A mutant variant which showed reduced vascular leak in vitro and reduced lethality in mice. In a mouse model of melanoma, we found significant decrease in tumor growth associated with reduction in Tregs when the protein was tested as monotherapy or in combination with checkpoint blockade.

## Introduction

The recent success of cancer immunotherapy epitomized by immune checkpoint blockade has contributed to a paradigm shift in cancer treatment. However, not all patients respond well to immune checkpoint blockade therapy, prompting the continued need for developing and improving novel therapies, including combination immunotherapies. The failure to induce robust anti-tumor immune responses in some checkpoint inhibitor recipients is likely due to alternative mechanisms of tumor-induced immune suppression such as the recruitment of tolerizing regulatory T cells (Tregs). Increased numbers of Tregs, correlating with poor prognosis, have been identified in many human cancers (1, 2), and studies in mice show that depletion of Tregs greatly increases the efficacy of immunotherapy (3–5). Hence, depletion of Tregs is a promising strategy for the enhancement of tumor-specific immunity.

Denileukin diftitox (DAB_1-389_IL-2, Ontak^®^) is a fusion protein toxin in which human IL-2 (hIL-2) is genetically linked to the N-terminal 389 amino acid fragment of diphtheria toxin (6). In this construct, the hIL-2 portion of denileukin diftitox replaces the native diphtheria toxin receptor binding domain and serves to target the cytotoxic action of the fusion protein to only those eukaryotic cells that display the high and intermediate affinity receptors for IL-2 (7). Once bound to the IL-2 receptor (IL-2R), denileukin diftitox is internalized by receptor mediated endocytosis and upon acidification of the vesicle lumen the translocation domain of the toxin partially denatures and spontaneously inserts into the membrane forming an 18 to 22Å pore (8–10). The catalytic domain of the fusion protein toxin is then thread through this pore, and its delivery and release into the cytosol is facilitated by COPI complex machinery and thioredoxin reductase (11, 12). Yamazumi *et al*. (13) demonstrated that the delivery of a single molecule of the catalytic domain into the eukaryotic cell cytosol is sufficient to kill that cell by the NAD^+^ dependent ADP-ribosylation of elongation factor 2 (EF-2), which halts protein synthesis.

In 1999, denileukin diftitox was approved by the Food and Drug Administration for the treatment of persistent cutaneous T cell lymphoma. Since the cytotoxic action of the fusion protein toxin is targeted toward only those eukaryotic cells that display the high and intermediate affinity IL-2R, the drug has been also used to eliminate CD25^+^ lymphoma cells (14), as well as T regulatory (Treg) and activated T effector (Teff) cells in syndromes ranging from stage IV unresectable malignant melanoma (15) to steroid resistant graft-versus-host disease (16).

Denileukin diftitox was produced commercially as a recombinant protein in *Escherichia coli* and expressed in high yield in the cytosol in inclusion bodies. In order to produce biologically active drug, partially purified inclusion bodies had to be completely denatured and then refolded in the presence Tween 20. Based on the observations of Boquet *et al*. (8), it is likely that Tween 20 was required in the refolding of denileukin diftitox to partially block intermolecular hydrophobic interactions of the toxin’s translocation domain that led to aggregation. Despite its clinical effectiveness (17), denileukin diftitox was placed on clinical hold in 2011 because of the presence of drug aggregates, contaminating DNA, varying concentrations of Tween 20, and batch to batch variations in its final formulation. The major dose-limiting toxicity of denileukin diftitox was vascular leak syndrome apparently unrelated to its CD25-specific toxicity.

In this paper, in order to address the vascular leak syndrome associated with denileukin diftitox, we have introduced a mutation (V6A) in one of the three vascular leak syndrome (VLS) inducing motifs found in the diphtheria toxin portion of the fusion toxin protein (18) to generate s-DAB_1-386_IL-2(V6A). Our data reveal that this mutation mitigates the vascular leak activity *in vitro* and reduces toxicity *in vivo,* allowing administration of higher doses.

Additionally, we completely reformulated the fusion toxin in order to produce disaggregated protein. Native diphtheria toxin is produced by toxigenic strains of *Corynebacterium diphtheriae* as a fully-folded soluble monomeric protein that is secreted into the culture medium. It is well known that the *tox* gene is carried by a family of closely related corynebacteriophages, and that regulation of *tox* gene expression is controlled by a *C. diphtheriae* host iron-activated repressor, DtxR (19). The production of diphtheria toxin requires growth of toxigenic strains of *C. diphtheriae* in low iron medium, and maximal yields are only obtained during the decline phase of growth when iron becomes the growth rate limiting substrate (20).

In order to circumvent the problems associated with cytosolic expression and refolding of denileukin diftitox, we used an *E. coli* / *C. diphtheriae* shuttle vector in the reconstruction of the structural gene for denileukin diftitox to direct its constitutive expression and secretion into the culture medium when grown in a non-toxigenic strain of *C. diphtheriae*. In this construct, we used the native *tox* promoter and a mutant *tox* operator to allow constitutive expression in iron rich medium, and the native *tox* signal sequence to direct the secretion of a denileukin diftitoxlike protein called s-DAB_1-386_IL-2, into the culture medium from which it can be readily purified in fully monomeric form. Finally, we demonstrate that s-DAB_1-386_IL-2(V6A) is highly potent both *in vitro* and *in vivo* in the treatment of B16 murine model of malignant melanoma as a monotherapy, as well as in a sequential immunotherapeutic regimen using s-DAB_1-386_IL-2(V6A) in combination with anti-murine-PD-1.

## Results

### Secretion of a denileukin-diftitox-like protein from *C. diphtheriae* enables simplified production of highly purified, monomeric fusion toxin from culture supernatants

After denileukin diftitox was placed on clinical hold in 2011, all available sources of the drug for both experimental and human use quickly became exhausted. In order to resolve issues that led to the drug being withdrawn from clinical use, principally to avoid aggregate contaminants that were found in the previous formulation of denileukin diftitox, we sought to develop a fusion toxin production method that did not require refolding of denatured protein or use of detergent (**Fig 1A**). During the early stages of the development of denileukin diftitox, the drug was in fact expressed as a secreted protein in the periplasmic space of recombinant *E. coli*, and purified from the periplasmic space as a fully biologically active IL-2R targeted toxin (IC_50_≥ 10^-11^M) by immunoaffinity chromatography using immobilized anti-diphtheria toxoid antibodies (21), although the yields were low.

**Fig. 1.**
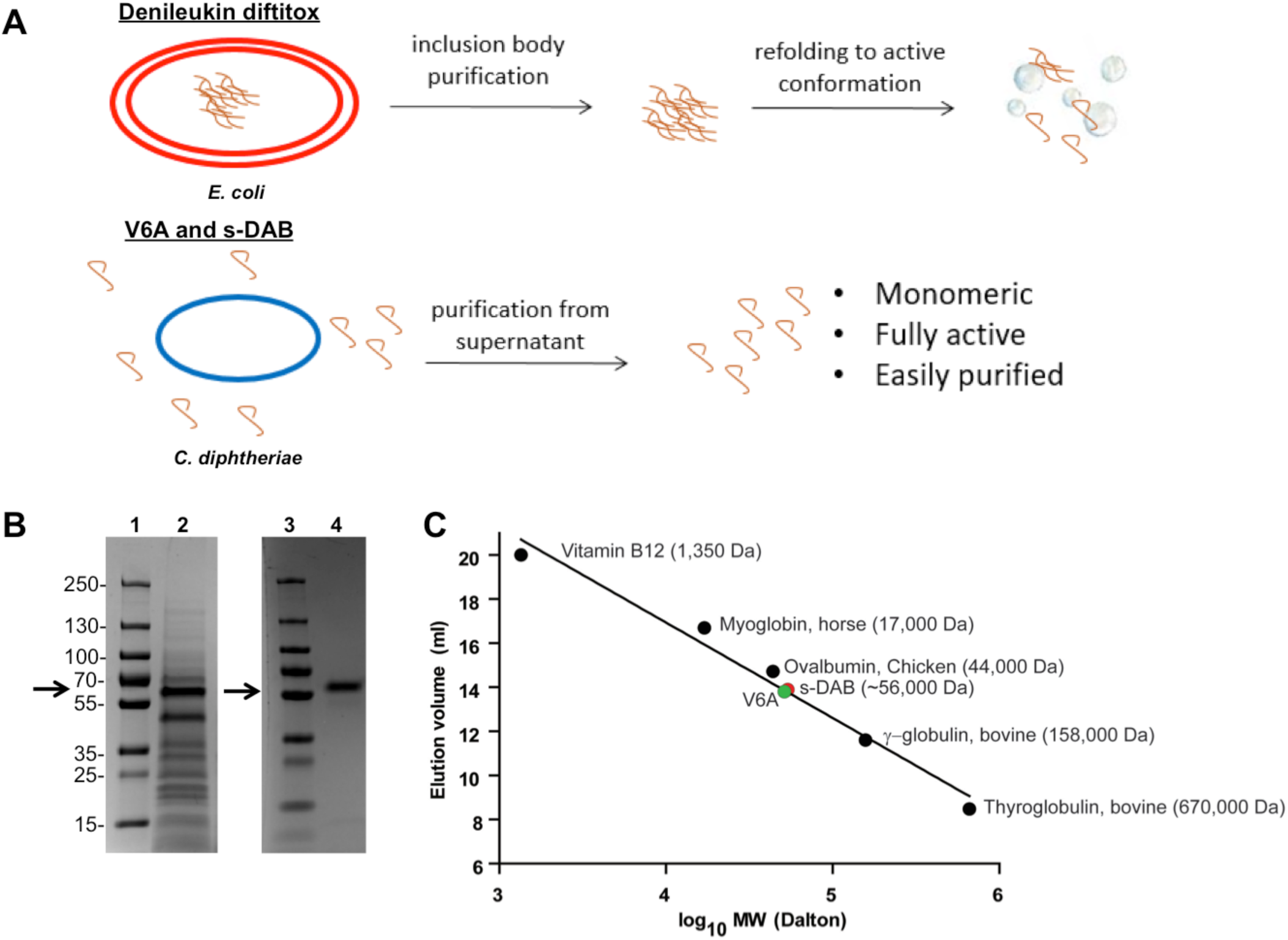
Production of s-DAB or V6A from *C. diphtheriae* generates monomeric and highly purified fusion toxin. (**A**) Schematic of fusion toxin production from *E. coli* versus *C. diphtheriae*. (**B**) Supernatant harvested from fermenter grown C7_s_(-)^tox-^ secreting V6A was analyzed by Coomassie-stained SDS-PAGE gel before (lane 2) and after purification (lane 4). Lanes 1 and 3 are molecular weight marker. Arrow indicates expected size of V6A at 56 kDa. (**C)** Elution profile of S-100 Superdex gel permeation chromatographic analysis of s-DAB, V6A and proteins of known molecular weight.

These prior observations encouraged us to re-construct the structural gene encoding denileukin diftitox with the native diphtheria *tox* signal sequence in order to direct the secretion of the fusion protein toxin (**Fig. 1*A*** and *SI Appendix,* **Fig. S1**). Further, since native diphtheria toxin is expressed in high yield from toxigenic strains of *C. diphtheriae*, we re-cloned the gene into an *E. coli* / *C. diphtheriae* shuttle vector pKN2.6Z, a Kn^R^ derivative of pCM2.6 (22). A transcriptional repressor, DtxR, binds the operator in the presence of iron, and repression is relieved under iron-limiting conditions. As iron is required for bacterial growth, diphtheria toxin is only expressed once iron is depleted (20). Therefore to increase the yield of mature DAB_1-_ 386IL-2 expressed as a secreted protein from the recombinant C7_s_(-)^tox-^ strain of *C. diphtheriae,* we used the native *tox* promoter and a mutant *tox* operator to allow constitutive expression in iron rich medium (*SI Appendix,* **Fig. S1)** (23, 24). Due to cleavage of the signal sequence during secretion, the diphtheria toxin portion of the fusion protein toxin is three amino acids shorter than denileukin diftitox. The final expression shuttle plasmid pKN2.6Z-LC127 (*SI Appendix,* **Fig. S2**) also included an origin of replication for both *E. coli* and *C. diphtheriae* and a C-terminal histidine tag to produce the secreted, His-tagged version of denileukin diftitox called s-DAB_386_-IL-2 (hereafter denoted as s-DAB).

We transformed pKN2.6Z-LC127 into the non-lysogenic non-toxigenic *C. diphtheriae* strain C7_s_(-)^tox-^, which was then grown in a fermenter, and the secreted protein was concentrated by tangential flow filtration using 30 kDa membrane followed by buffer exchange by diafiltration. The concentrate was affinity purified using a Ni^2+^ affinity column, and subsequently purified by an S-100 gel permeation sizing column, which allowed separation of full length s-DAB from smaller proteolytically degraded protein fragments. The resulting product was ∼95% pure as shown by Coomassie blue-stained SDS-PAGE gel analysis (**Fig. 1*B***) with a yield of 809 µg s-DAB produced per liter of culture. It was found to be a monomeric 56 kDa protein by size-exclusion chromatography (**Fig. 1*C***).

### V6A substitution in denileukin diftitox leads to markedly reduced vascular leak

Approximately 25% of patients treated with denileukin diftitox developed vascular leak syndrome (VLS), and it was a common dose-limiting toxicity (17). VLS results from damage to vascular endothelial cells leading to fluid extravasation into tissues. Although transient at times, VLS can be severe, leading to hypotension, hypovolemia, peripheral edema, pleural and pericardial effusions, and in rare cases, multi-organ dysfunction. The molecular mechanism of VLS is not well understood. Natural killer (NK) cells can target endothelial cells for lysis, and in mice lacking NK cells, IL-2-induced vascular leak was significantly reduced (25). Inflammatory cytokines TNFα, IL-1, and IL-2 have also been shown to cause vascular leak in animal models (26–28).

Baluna and colleagues (18) showed that the (x)D(y) motif (where x = L, I, G or V and Y= V, L, or S) in ricin toxin disrupted the integrity of HUVEC monolayers. They demonstrated that mutations in these motifs in ricin led to decreased disruption of endothelial cell monolayers *in vitro*, and decreased the induction of vascular leak in mice. They reported the presence of several VLS-inducing motifs in hIL-2 and diphtheria toxin sequences. Since denileukin diftitox and s-DAB have VLS motifs, we hypothesized that mutating these motifs may mitigate the adverse effects of the drug. We therefore used site-directed mutagenesis to make a single V6A amino acid substitution in s-DAB. Since the N-terminal predicted VLS-inducing motif of s-DAB (residues 6-8) is not part of the ADP-ribosyltransferase active site, we anticipated this substitution would not affect the protein’s catalytic activity. The VDS motif is also located in an exposed random coil in the N-terminal region of the fusion protein, and therefore was most likely to affect cell-cell and, or cell-matrix interactions (29).

We found that the expression of His-tagged s-DAB_1-386_IL-2(V6A) (hereafter denoted as V6A) from the native *tox* promoter / mutant *tox* operator was constitutive. Following purification as standardized for s-DAB, the V6A mutant eluted from an S-100 column in a single ≥ 97% homogeneous peak corresponding to an V_e_ of 56 kDa, showing that it was also secreted into the culture medium as a fully-folded monomeric recombinant protein (**Fig. 1*C***).

### V6A substitution retains biologic activity, but significantly reduces vascular leak

To assess whether the V6A mutation affects activity, we performed a dose response analysis of s-DAB and V6A in an *in vitro* cytotoxicity assay against MT-2 cells, which express CD25 (7) and are sensitive to killing by denileukin diftitox. We found that s-DAB and V6A have IC_50_ values of 0.12 and 0.33 pM, respectively, suggesting that the V6A mutation did not significantly affect biologic activity (**Fig. 2*A***). The reported IC_50_ for classic denileukin diftitox, is 2 pM in the closely related HUT-102 cell line (7), which is substantially higher than that for either s-DAB or V6A indicating that a significant enhancement of potency with the monomeric rather than aggregated versions of the fusion proteins.

**Fig. 2.**
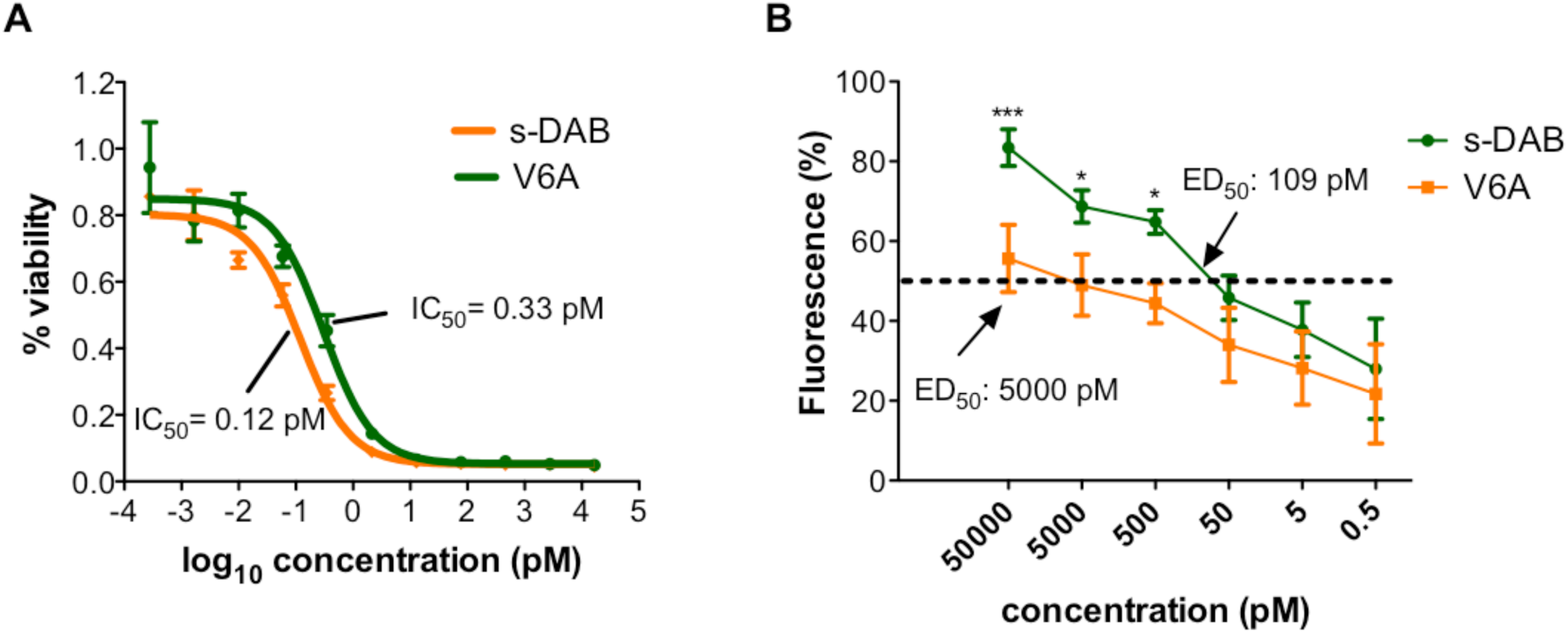
V6A exhibits comparable cytotoxic activity and induces less vascular permeability than s-DAB *in vitro*. (**A**) Activity of V6A and s-DAB against the CD25+ cell line MT-2 *(6)* in an MTS cell killing assay showed an IC_50_ of 0.33 and 0.12 pM for V6A and s-DAB, respectively. (**B**) FITC-labeled dextran permeation assay in HUVEC cells. Cells were seeded in the upper well of a Transwell chamber and treated with V6A, s-DAB, or LPS for 19 hrs. The amount of FITC-dextran which permeates to the lower well was measured by fluorescence. Values represent percent of fluorescence of LPS positive control treated cells. Three independent experiments were performed in duplicate. Statistical analysis of the results was carried out using ANOVA. Error bars show SEM. ***p < 0.001 and *p < 0.05.

We then tested s-DAB and V6A in a HUVEC permeability assay in order to assess their ability to cause vascular leak. When tested at concentrations ranging from 0.5 pM to 50 nM, we found that the V6A mutant showed statistically significant reduction in permeability compared to s-DAB at 500 pM, 5 nM and 50 nM (**Fig. 2*B***). Indeed, the ED_50_ for vascular permeation for V6A was 5 nM while that for s-DAB was 109 pM revealing an approximately 50-fold reduction in vascular leak with V6A compared to s-DAB.

We also studied the toxicity of V6A versus s-DAB in a mouse chronic administration study giving each drug at 1, 3.2, 10, and 32 µg (the equivalent of 0.05, 0.16, 0.5, and 1.6 mg/kg, respectively) daily for 15 days. Mice receiving doses of 3.2 µg or higher of s-DAB lost weight for the duration of the experiment, while the weights of mice given 3.2 µg V6A were indistinguishable from control mice (**Fig. 3*A***). Additionally, all mice receiving 10 µg of s-DAB daily died with 8 days, while there was no lethality in mice receiving V6A 10 µg daily (Fig. 3B). The calculated LD_50_ for V6A was 18.2 µg daily (0.91 mg/kg) while it was 4.9 µg daily (0.24 mg/kg) for s-DAB (30). Thus, the LD_50_ of V6A is 50-fold higher than the maximal clinically recommended dose of denileukin diftitox (18 µg/kg daily for 5 days) and is 4.1-fold higher if adjusted for allometric scaling based on body surface area and murine metabolism (31).

**Fig. 3.**
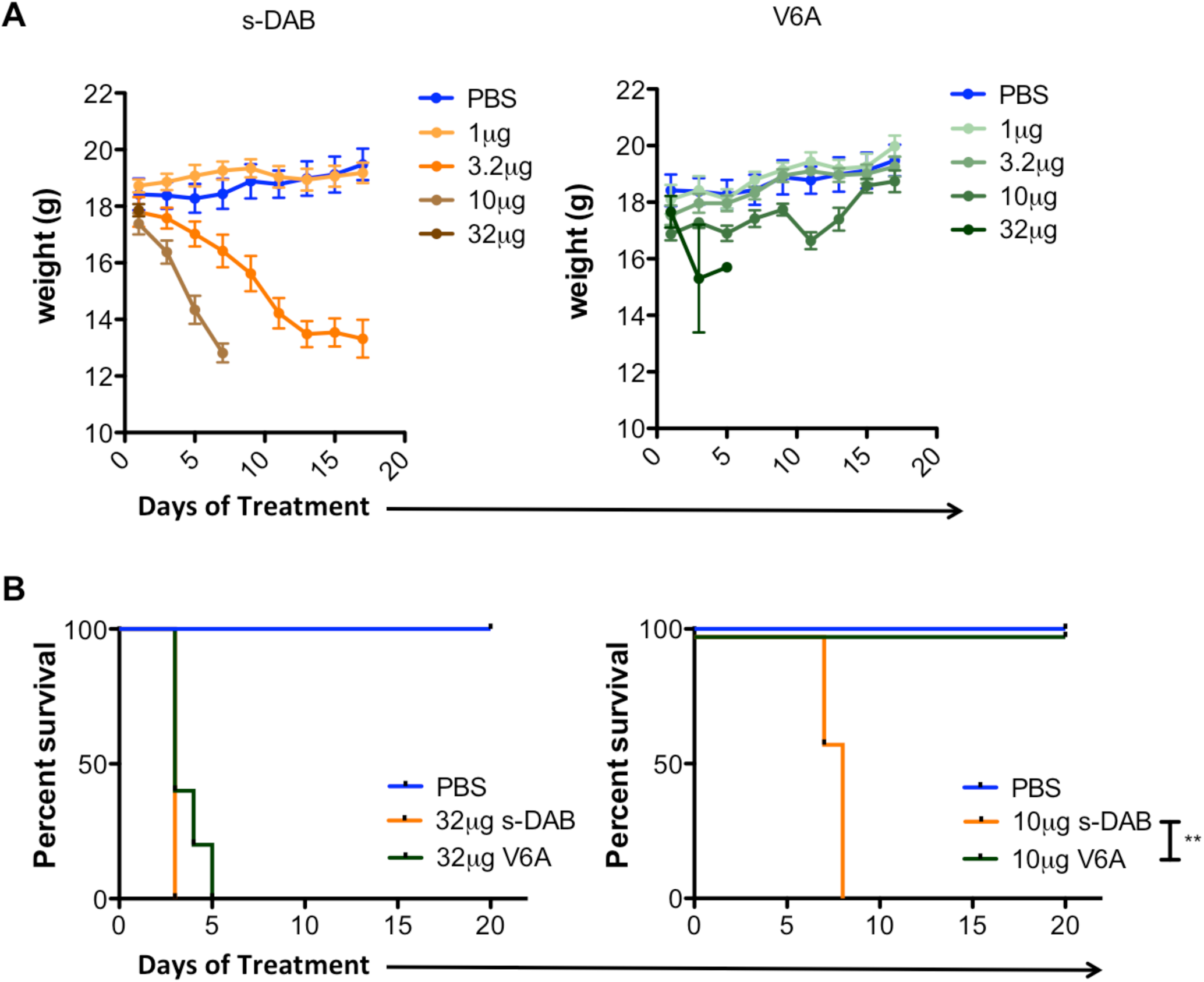
V6A treatment induces less toxicity than s-DAB in mice. Mice were treated daily with PBS, s-DAB, or V6A at indicated doses for 20 days (n=5). (**A)** Weights of treated mice. Error bars are mean ± SEM. (**B**) Survival of treated mice. Statistical analysis by log-rank test. **p<0.01.

It has been shown that protein aggregates exacerbate infusion reactions (32), so classic denileukin diftitox which contains 40% aggregates, and is contaminated by detergent, is likely to cause more vascular leak than s-DAB, which is monomeric and has no contaminating detergent. However, we could not test this directly since classic denileukin diftitox is no longer available. Nevertheless, our data clearly reveal significantly reduced vascular leak, toxicity, and lethality with the V6A version of s-DAB.

### V6A depletes Treg cells, is active in tumor models, and potentiates the activity of PD-1 blockade

We examined the effectiveness of V6A both as a monotherapy and in a sequential immunotherapeutic regimen using anti-PD-1 antibody in the B16F10 murine model of malignant melanoma, in which mice develop poorly immunogenic tumors that are highly infiltrated with Tregs (33). As denileukin diftitox is known to transiently deplete Tregs *in vivo*, we compared lymphocyte populations in melanoma-bearing mice treated with s-DAB or V6A. Mice were treated on days 7 and 10 post B16F10 cell implantation. Both drugs depleted Tregs in the lymph nodes and spleens of tumor bearing mice with equivalent efficacy (**Fig. 4*A***). s-DAB and V6A treament also significantly inhibited tumor growth with similar efficacy (**Fig. 4*B***), suggesting that the V6A mutant could be as effective in promoting anti-tumor responses as denileukin diftitox without the reported vascular leak side effects.

**Fig. 4.**
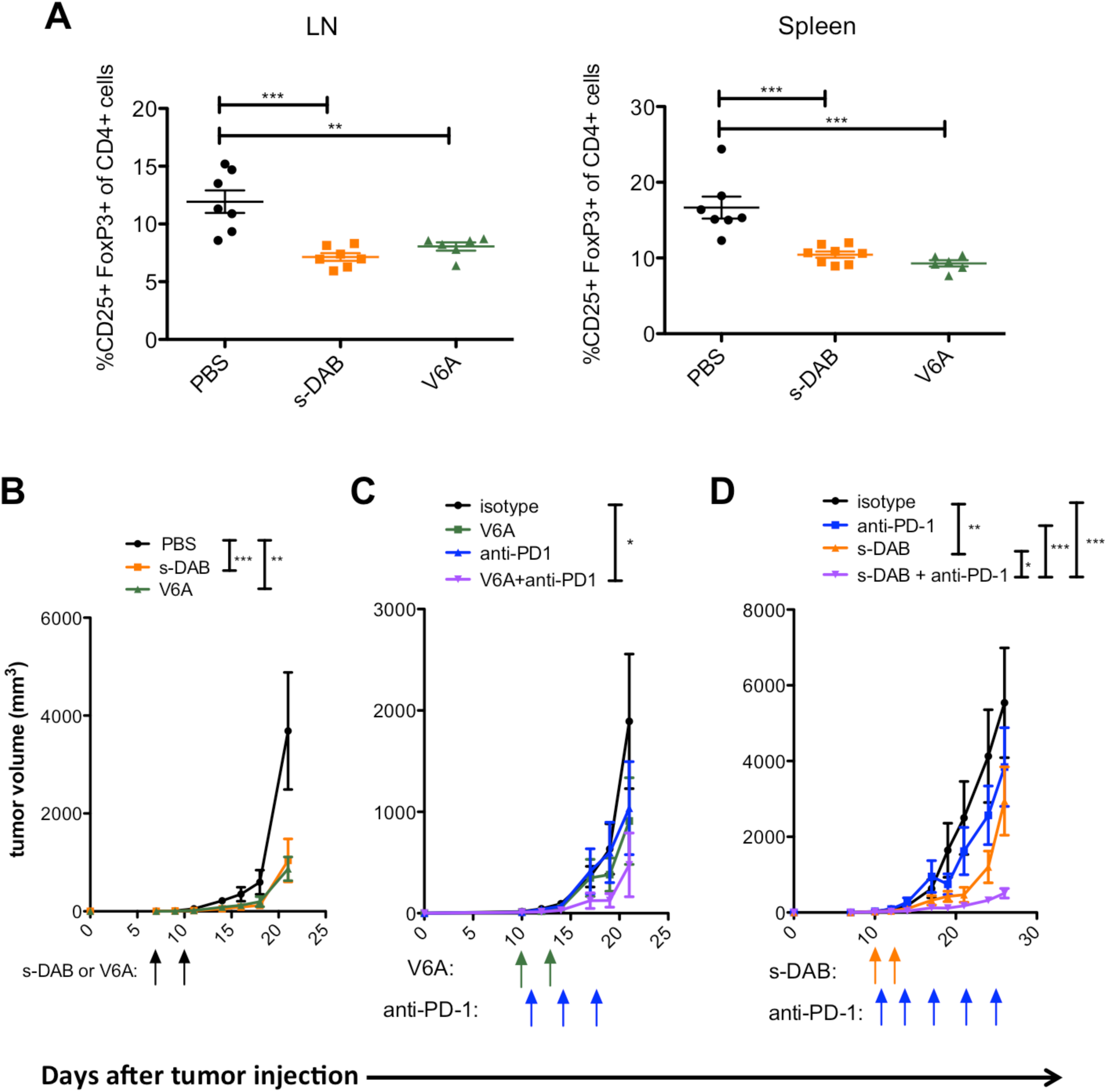
Anti-tumor activity of V6A is equivalent to s-DAB and enhances the efficacy of anti-PD-1 in the B16F10 melanoma model. **(A-B)** C57BL/6 mice were treated with 5 µg of s-DAB or V6A on day 7 and day 10 post-B16F10 inoculation. (**A**) Percentage of CD25+FoxP3+ Tregs of CD4+ T cells in tumor-draining lymph nodes (LN) and spleens at day 24 post B16F10 inoculation Statistical significance was measured by one-way ANOVA. (n=6-8). (**B**) Tumor size was measured by electronic caliper (n=8). Statistical significance for tumor growth experiments was assessed by repeated measures two-way ANOVA on day 18 (PBS vs s-DAB, p<.001; PBS vs V6A, p<0.01). (**C**) C57BL/6 mice were treated with 10 µg V6A on day 10 and day 13 post-B16F10 inoculation. Anti-PD-1 or isotype control was begun on day 11 and given twice per week for the duration of the experiment (n=8) (isotype vs V6A+anti-PD-1, p<0.05). (**D**) C57BL/6 mice were treated with 5 µg s-DAB on days 10 and 13 post B16F10 inoculation. Anti-PD-1 or isotype control was begun on day 11 post B16F10 inoculation and given 2 times per week for the duration of the experiment. (isotype vs s-DAB, p<0.01; isotype vs anti-PD-1, anti-PD-1 vs s-DAB + anti-PD-1, p<0.001; s-DAB vs s-DAB + anti-PD-1, p<0.05). Tumor size was measured by electronic caliper. Arrows are days treatments were given; green: V6A; blue: anti-PD-1; orange: s-DAB. Data are shown as mean ± SEM. Statistical significance for tumor growth experiments were assessed by repeated measures two-way ANOVA with Bonferroni post-test for multiple comparisons * p<.05, **p<0.01, *** p<0.001

Programmed cell death-1 (PD-1) and its receptor programmed cell death ligand-1 (PDL-1) are key regulatory checkpoints that maintain self-tolerance and the degree of T and B cell activation (34, 35). Many types of cancers over-express PD-L1, which then binds to PD-1 on T effector cells infiltrating the tumor, inhibiting their anti-tumor activity. This immune checkpoint circuit has become a major target in the development of immunotherapeutic agents, with remarkable clinical responses seen in a subset of cancer patients (36, 37). We therefore reasoned that the sequential use of V6A followed by an anti-PD-1 regimen might transiently deplete CD25^+^ FoxP3^+^ Treg suppression of an anti-tumor response followed by stimulation of a T effector anti-tumor response, thereby promoting an enhanced immune response against the tumor.

In order to better observe the effects of dual therapy on tumor growth, tumors were allowed to progress for a longer time before treatment initiation. Melanoma-bearing mice were treated 10 days after tumor initiation with PBS, an isotype control, anti-PD-1, s-DAB or V6A as monotherapies. Mice were treated on days 10 and 13 with s-DAB or V6A, and with anti-PD-1 on days 11, 14, and twice weekly thereafter. When used as monotherapies in more progressed tumors, anti-PD-1, s-DAB and V6A had no effect on tumors when compared to mice that were treated with an isotype control antibody (**Fig. 4 *C* and *D*)**. In contrast, when administered as sequential immunotherapies, V6A or s-DAB in conjunction with anti-PD-1 elicits a remarkable inhibition of tumor growth (**Fig. 4 *C* and *D***).

Treg cells isolated from metastatic lymph nodes from melanoma are known to inhibit CD4^+^ and CD8^+^ T-cell proliferation *in vitro*. To examine if the reductions in Treg levels and decreased tumor growth observed in mice treated with s-DAB or V6A are associated with improved anti-tumor immunity in the tumor microenvironment, we excised tumors from surviving mice in each group and analyzed tumor infiltrating lymphocytes for IFNγ production. We found a significant increase in IFNγ production by CD8^+^ cytotoxic T cells in spleens and tumors from mice that were treated with s-DAB and observed a similar trend in V6A treated mice (**Fig. 5**).

**Fig. 5.**
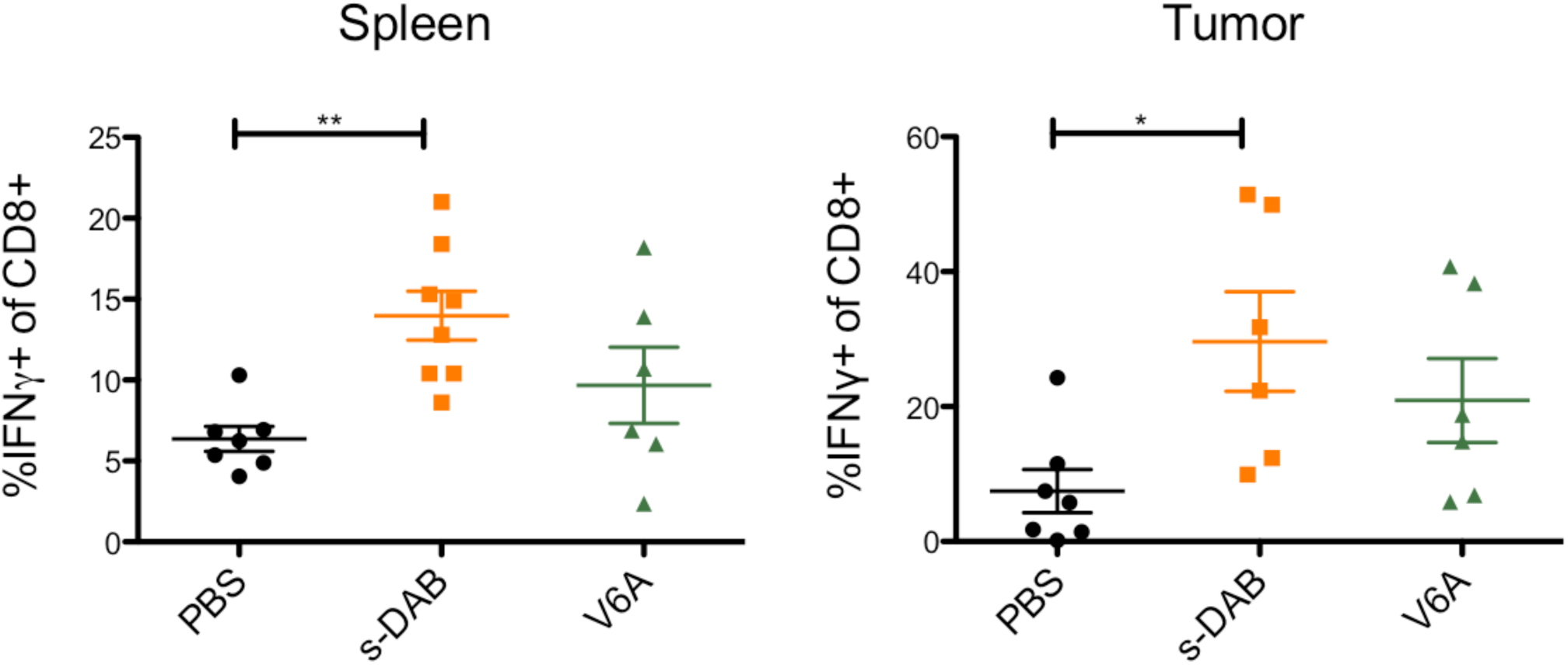
Effects of fusion toxin treatment on CD8+ lymphocytes in tumors and spleens. Mice were treated with 5 µg s-DAB or V6A on days 7 and 10 post B16F10 inoculation (n=6-8). Tumors and spleens were harvested on day 24 and analyzed by flow cytometry. Statistical significance determined by one-way ANOVA. * p<.05, **p<0.01.

## Discussion

We report the development of a second-generation version of denileukin diftitox fusion protein called s-DAB_1-386_IL-2(V6A). By re-designing production as a secreted fully-folded monomeric fusion protein toxin in *C. diphtheriae*, V6A overcomes the production problems of inactive aggregates and contaminating detergent that plagued the refolding of denileukin diftitox from denatured inclusion bodies. Indeed our results show that, in addition to being fully monomeric, V6A is more potent *in vitro* (IC_50_ for CD25+ MT-2 cells of 0.33 pM) than the reported values for denileukin diftitox (2 pM) most likely because it lacks high levels of inactive aggregates. Moreover, we re-engineered V6A to reduce vascular leak, which was a serious, sometimes treatment-limiting adverse effect of denileukin diftitox. By substituting Val-6 with Ala, V6A retains potent killing activity against CD25+ cells, but it elicits 50-fold less vascular leak *in vitro* than its parent protein also made in *C. diphtheriae*, s-DAB. V6A is better tolerated in mice receiving high doses than is s-DAB, and it has a mouse LD_50_ of 0.910 mg/kg which is 3.7-fold higher than that of s-DAB.

Our work also demonstrates for the first time that Treg-depleting fusion toxins based on denileukin diftitox show potent enhancement of checkpoint inhibitor therapy. Indeed, in more progressed tumors that were unresponsive to single agent therapy, the addition of fusion proteins to anti-PD-1 therapy resulted in significant anti-tumor efficacy. While classic denileukin diftitox was previously studied in the mouse B16 melanoma model in combination with anti-PD-L1 and anti-CTLA4, no additive effects were seen (38). Our positive findings are likely due to the lower potency of denileukin diftitox (due to inactive aggregates) compared with V6A and s-DAB as well as use of a higher dose of fusion toxin in our studies. Additionally, in our work, tumors were allowed to progress for 10 days before the first administration of fusion toxin, while earlier studies begin treatment at earlier time points (38, 39). The fact that we observed antitumor efficacy even with treatment initiation at a later time point underscores the greater potency of our fully monomeric, *C. diphtheriae*-derived fusion proteins.

Being highly immunogenic among the common adult-onset cancers, melanoma is believed to be the most responsive to intervention by immunotherapy. Although denileukin diftitox was FDA approved for the treatment of patients with cutaneous T-cell lymphoma (CTCL), CTCL cells share many features with Treg cells, including expression of CTLA-4, Foxp3, and suppression of CD4^+^ T-cell activation. Treatment of mice with anti-CD25 antibody has been found to promote CD8+ T cell-mediated tumor immunity through depletion of Tregs (40). CD4^+^CD25^hi^ Treg cells are also reported to be increased in the lymph nodes of patients with metastasic melanoma (41). In a Phase II trial of denileukin diftitox in patients with unresectable stage IV melanoma, 45.5% of immune- and chemotherapy naïve patients who were treated achieved at least a partial response, and one year later their survival rate was markedly higher than patients who did not receive the drug (42). These findings encouraged us to examine the effectiveness of s-DAB and V6A in the B16 murine model of malignant melanoma.

Blocking immune checkpoint molecules such as CTLA-4 and PD-1 has provided clinical success in malignant melanomas and lung cancers by allowing a resurgence in the effector function of tumor-infiltrating T cells. Yet, more than half of the treated patients do not respond to immune checkpoint blockade therapy. Depletion of Tregs using anti-CD25 antibodies has been shown to enhance anti-tumor efficacy of anti-PD-1 in mice (43). However, activated effector T cells also express CD25, and the use of anti-CD25-antibodies (which have long serum half-lives) for Treg depletion may concomitantly reduce activated effector T cells as well thereby countering any beneficial effect of Treg depletion to augment anti-tumor immunity (44).

Since V6A and s-DAB deplete Tregs transiently (similar to denileukin diftitox), with rebound observed by 72 hours post treatment (15), side effects due to hyperinflammation are not expected. Based upon classic denileukin diftitox, we have taken advantage of the anticipated short half-life (*i.e*., 72 minutes) (45) of s-DAB and V6A to promote initial depletion of Tregs in a melanoma model, and then stimulate the T effector arm of the immune system with anti-PD-1. Using this sequential approach to combination therapy, we found significant decreases in tumor growth associated with reduction of Tregs and enhancement of effector T cells. Thus s-DAB_1-_ 386IL-2(V6A) is a next generation of denileukin diftitox that shows potential for improved clinical utility in patients warranting further investigation as a cancer immunotherapeutic, both as a monotherapy and in combination with immune checkpoint blockade.

## Materials and Methods

### Plasmids, bacterial strains, and cloning

pKN2.6z *E. coli*-*C. diphtheriae* shuttle vector was kindly provided by Dr. Michael P. Schmitt (Food and Drug Administration, Silver Spring MD). Plasmids transformed by into either chemically competent NEB 5-alpha *E. coli* (NEB) or electrocompetent *C. diphtheriae* C7_s_(-)^tox-^. Fusion toxin constructs were synthesized and cloned into pKN2.6z vector using NEBuilder HiFi Assembly Cloning Kit (NEB).

### Fermenter protein production and purification

C7_s_(-)^tox-^ was transformed with plasmids carrying the fusion toxin gene and were grown in CY medium overnight in a shaking incubator at 37°C (REF 103). The next day, 10 mL of shaking culture was used to inoculate 1.4 L of CY medium in a fermenter (New Brunswick, Bioflo/Celligen 110) and grown at 30°C, air at 2 L/min, agitation at 450 rpm for 12 hours and then increased to 36°C. Optical density (OD) at 600 nm was measured hourly. Bacteria were pelleted by centrifugation and supernatant was harvested once the OD reached 12-15. Supernatant was concentrated 20-fold by tangential flow filtration and through a 30kD hollow fiber membrane (Spectrum) and then diafiltered against 2 L of 50 mM NaH_2_PO_4_, 500 mM NaCl, 50 mM imidazole pH7.4. Concentrate was applied to a HisTrap HP column (GE Healthcare) and 6x His tagged protein was eluted with 50 mM NaH_2_PO_4_, 500 mM NaCl, 500 mM imidazole pH7.4. Eluent was concentrated to 5 mL with 10 kDa Amicon Ultra-15 centrifugal units (Millipore Sigma) and run over a HiPrep 26/60 Sephacryl S100-HR sizing column (GE Healthcare Life Sciences) eluted with PBS. 5 mL fractions were collected and analyzed by SDS-PAGE and ImageJ software. Protein was stored at −80°C in 15% glycerol and PBS.

### In vitro cytotoxicity assays

The following reagent was obtained through the AIDS Reagent Program, Division of AIDS, NIAID, NIH: MT-2 cell line from Dr. John Riggs. Cell lines were grown in 90% RPMI-1640 supplemented with 10% heat-inactivated fetal bovine serum (Gibco). s-DAB and V6A was tested for cytotoxicity against MT-2 cells. Briefly, cells were plated at a density of 1 x 10^6^ cells per mL in triplicate and treated with 10-fold serial dilutions of fusion toxin or no drug. Cells were incubated for 48 hrs with fusion toxin and MTS reagent (Promega) was used to measure proliferation. Absorbance was read at 490 nM with the iMark Microplate Reader (Bio-Rad).

### In vitro vascular permeability assay

4X10^5^ HUVEC cells were seeded on Transwell collagen-coated, 0.4µm pore polyethylene membrane inserts in 24-well plates and cultured with 250 µl of the culture medium in the upper chamber and 500 µl of the same growth medium in the lower chamber. Cells were grown for three days without changing the medium until they reached confluence. Cells were treated with various concentration of V6A, s-DAB, and LPS for 19 hrs. Fluorescein isothiocyanate (FITC)-conjugated dextran (molecular weight 40 000) was added to the upper chamber of all wells. After a 30-minute incubation at 37°C, the bottom well media were transferred into a 96-well black plate, and fluorescence was measured using a spectrofluorimeter (excitation, 485 nm; emission, 535 nm). Fluorescence represents the amount of FITC-dextran passing from the top chamber to the bottom chamber.

### Mouse toxicity and LD_50_ determination

C57BL/6 mice were purchased from the Charles Rivers Laboratory and animal studies were performed according to IACUC approved protocols at Johns Hopkins University. To assess toxicity, s-DAB, V6A, or PBS was given by intraperitoneal injection daily at specified doses. LD_50_ was determined by Reed-Meunch analysis (30).

### B16F10 melanoma tumor model and treatments

C57BL/6 mice were administered s-DAB or V6A by intraperitoneal injection on specified days. For melanoma experiments, mice were given subcutaneous injections of 1 x 10^5^ B16F10 cells in the right flank. 100 µg of anti-PD-1 (Bio X Cell) was given by intraperitoneal injection twice per week. Tumors were measured by electronic caliper and tumor volume was calculated using the following equation: tumor volume = length x width x height 0.5326. Mice were sacrificed at specified time points and tumor draining lymph nodes, spleens and tumors were isolated. Single-cell suspensions were prepared by dissociation through 100 µm filters. Red blood cells from spleens were lysed in ACK lysing buffer. Tumor-infiltrating lymphocytes were isolated by density gradient centrifugation in 80%/40% Percoll (GE Healthcare).

### Flow cytometry analysis

Single-cell suspensions were stained for viability using the LIVE/DEAD Fixable Aqua Dead Cell Stain Kit (Thermo Fisher Scientific). Cells were incubated with purified rat anti-mouse CD16/32 (BD) and labeled in FACS buffer (PBS, 2% heat-inactivated FBS, 0.1% HEPES, 0.1%) sodium azide) with the following antibodies (BD unless otherwise noted): Ax700 anti-CD8, APC anti-CD4, and BV421 CD25. For *in vitro* stimulations, cells were incubated with PMA (50 ng/mL) and ionomycin (1µM) with GolgiPlug (BD) for 4 hours at 37°C. Surface staining was performed as above, and intracellular staining was performed with the Fixation/Permeabilization Solution Kit (BD) according to manufacturer’s protocol and labeled with FITC IFNγ. Samples were acquired on an LSRII (BD) and data was analyzed using FlowJo (Tree Star).

## Acknowledgements

We are grateful to Michael Schmitt for advice, strains and plasmid pKN2.6Z, and to Brian Herbst for cloning and expression assistance.

## Author contributions

Conception and design: L.S.C, J.F., J.R.M, D.M.P, and W.R.B.; development of methodology: L.S.C, J.F, P.K, A.K, G.P., R.H., J.R.M, D.M.P, and W.R.B.; acquisition of data: L.S.C, J.F, P.K, A.K. M.E.U., E.A.I, C.K.B, G.P, J.R.M.; analysis and interpretation of data: L.S.C, J.F, P.K, A.K. M.E.U., E.A.I, S.P, C.K.B, G.P, R.H., J.R.M., D.M.P, and W.R.B; writing, review, and revision of manuscript: L.S.C, J.R.M, D.M.P, and W.R.B.

## Supplementary Information for

A second generation IL-2 receptor-targeted diphtheria fusion toxin exhibits anti-tumor activity and synergy with anti-PD-1 in melanoma

### This PDF file includes

Figs. S1 to S2

**Fig. S1.**
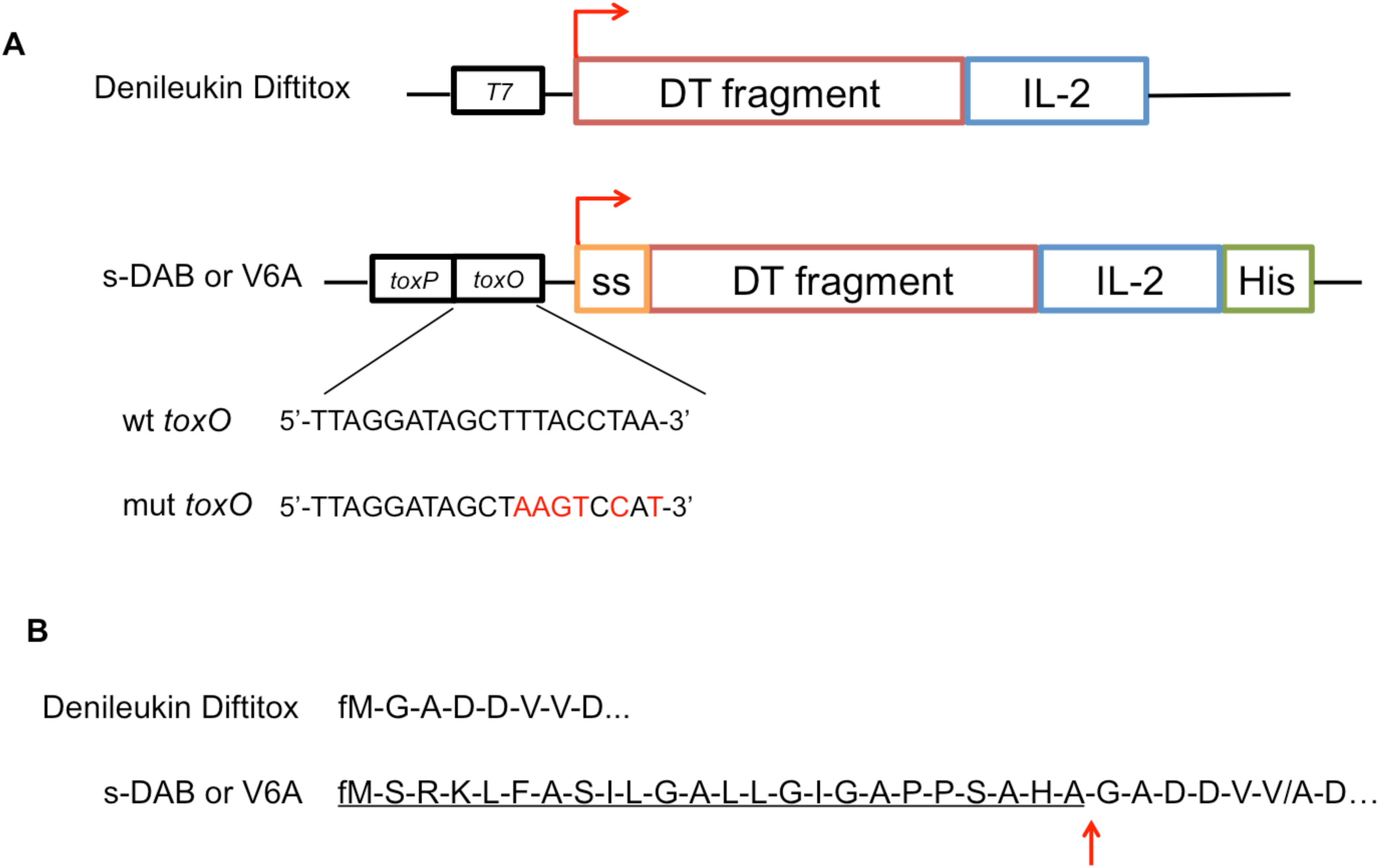
Genetic construct to secrete fusion toxin

**Fig. S2.**
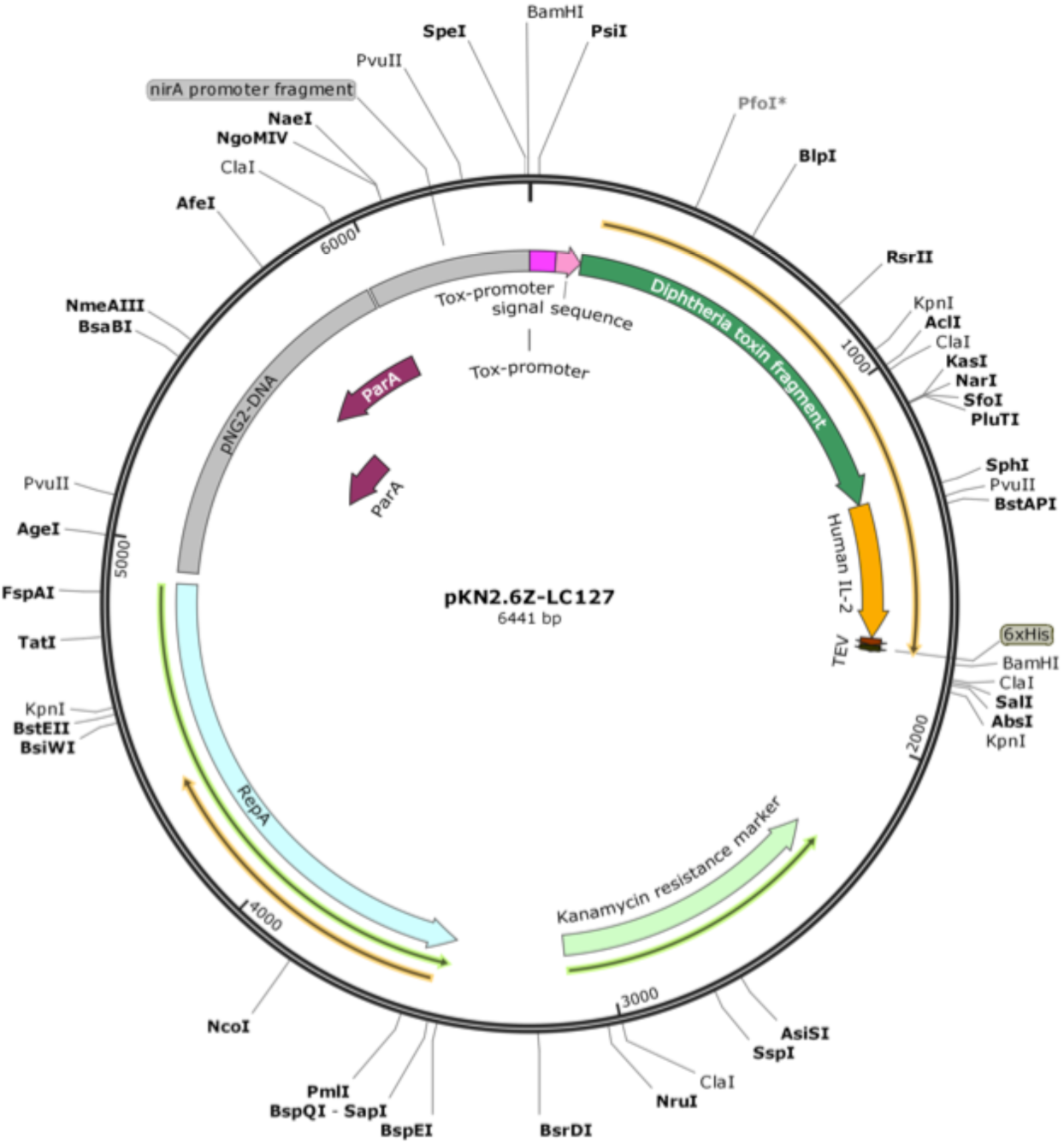
pKN2.6Z-LC127 shuttle vector plasmid map.

## References

1. Shang B, Liu Y, Jiang S, Liu Y (2015) Prognostic value of tumor-infiltrating FoxP3+ regulatory T cells in cancers: a systematic review and meta-analysis. Sci Rep 5(15179). doi:10.1038/srep15179.

2. Shah W, et al. (2011) A reversed CD4/CD8 ratio of tumor-infiltrating lymphocytes and a high percentage of CD4+ FOXP3+ regulatory T cells are significantly associated with clinical outcome in squamous cell carcinoma of the cervix. Cell Mol Immunol 8:59–66.

3. Shimizu J, Yamazaki S, Sakaguchi S (1999) Induction of tumor immunity by removing CD25+CD4+ T cells: a common basis between tumor immunity and autoimmunity. J Immunol 163:5211–5218.

4. Liu J, et al. (2016) Assessing immune-related adverse events of efficacious combination immunotherapies in preclinical models of cancer. Cancer Res 76(18):5288–5302.

5. Taylor NA, et al. (2017) Treg depletion potentiates checkpoint inhibition in claudin-low breast cancer. J Clin Invest 127(9):3472–3483.

6. Williams D, Snider CE, Strom TB, Murphy JR (1990) Structure/function analysis of interleukin-2-toxin (DAB486-IL-2). J Biol Chem 265(20):11885–11889.

7. Re GG, et al. (1996) Interleukin 2 (IL-2) receptor expression and sensitivity to diphtheria fusion toxin DAB389IL-2 in cultured hematopoietic cells. Cancer Res 56:2590–2595.

8. Boquet P, Silverman MS, AM Pappenheimer Jr., Vernon WB (1976) Binding of Triton X-100 to diphtheria toxin, crossreacting material 45, and their fragments. Proc Natl Acad Sci U S A 73(12):4449–4453.

9. Donovan JJ, Simon MI, Draper RK, Montal M (1981) Diphtheria toxin forms transmembrane channels in planar lipid bilayers. Proc Natl Acad Sci U S A 78(1):172–176.

10. Kagan BL, Finkelstein a, Colombini M (1981) Diphtheria toxin fragment forms large pores in phospholipid bilayer membranes. Proc Natl Acad Sci U S A 78(8):4950–4.

11. Ratts R, et al. (2003) The cytosolic entry of diphtheria toxin catalytic domain requires a host cell cytosolic translocation factor complex. J Cell Biol 160(7):1139–1150.

12. Trujillo C, Taylor-Parker J, Harrison R, Murphy JR (2010) Essential lysine residues within transmembrane helix 1 of diphtheria toxin facilitate COPI binding and catalytic domain entry. Mol Microbiol 76(4):1010–1019.

13. Yamaizumi M, Mekada E, Uchida T, Okada Y (1978) One molecule of diphtheria toxin fragment A introduced into a cell can kill the cell. Cell 15:245–250.

14. Lemaistre CF, et al. (1998) Phase I trial of a ligand fusion-protein (DAB 389 IL-2) in lymphomas expressing the receptor for interkeukin-2. Blood 91(2):399–406.

15. Rasku MA, et al. (2008) Transient T cell depletion causes regression of melanoma metastases. J Transl Med 6(12). doi:10.1186/1479-5876-6-12.

16. Ho VT, et al. (2004) Safety and efficacy of denileukin diftitox in patients with steroid-refractory acute graft-versus-host disease after allogeneic hematopoietic stem cell transplantation. Blood 104(4):1224–1226.

17. Olsen E, et al. (2001) Pivotal phase III trial of two dose Levels of denileukin diftitox for the treatment of cutaneous T-cell lymphoma. J Clin Oncol 19(2):376–388.

18. Baluna R, Rizo J, Gordon BE, Ghetie V, Vitetta ES (1999) Evidence for a structural motif in toxins and interleukin-2 that may be responsible for binding to endothelial cells and initiating vascular leak syndrome. Proc Natl Acad Sci U S A 96(7):3957–62.

19. Boyd J, Oza MN, Murphy JR (1990) Molecular cloning and DNA sequence analysis of a diphtheria tox iron-dependent regulatory element (dtxR) from Corynebacterium diphtheriae. Proc Natl Acad Sci U S A 87:5968–5972.

20. Pappenheimer AM (1977) Diphtheria toxin. Annu Rev Biochem 46:69–94.

21. Williams DP, et al. (1987) Diphtheria toxin receptor binding domain substitution with interleukin-2: genetic construction and properties of a diphtheria toxin-related interleukin-2 fusion protein. Protein Eng 1(6):493–498.

22. Drazek ES, Hammack CA, Schmitt MP (2000) Corynebacterium diphtheriae genes required for acquisition of iron from haemin and haemoglobin are homologous to ABC haemin transporters. Mol Microbiol 36(1):68–84.

23. Boyd J, Murphy JR (1988) Analysis of the diphtheria tox promoter by site-directed mutagenesis. J Bacteriol 170(12):5949–5952.

24. Tao X, Murphy JR (1994) Determination of the minimal essential nucleotide sequence for diphtheria tox repressor binding by in vitro affinity selection. Proc Natl Acad Sci U S A 91(20):9646–9650.

25. Assier E, et al. (2004) NK cells and polymorphonuclear neutrophils are both critical for IL-2-induced pulmonary vascular leak syndrome. J Immunol 172:7661–7668.

26. Goldblum SE, Hennig B, Jay M, Yoneda K, Mcclain CJ (1989) Tumor necrosis factor a-induced pulmonary vascular endothelial injury. Infect Immun 57(4):1218–1226.

27. Martin S, Maruta K, Burkart V, Gillis S, Kolb H (1988) IL-1 and IFN-gamma increase vascular permeability. Immunology 64(February):301–305.

28. Rafi-Janajreh AQ, et al. (1999) Evidence for the involvement of CD44 in endothelial cell injury and induction of vascular leak syndrome by IL-2. J Immunol 163(3):1619–27.

29. Bennet MJ, Choe S, Eisenberg D (1994) Refined structure of dimeric diphtheria toxin at 2.0 A resolution. Protein Sci 3:1444–1463.

30. Reed LJ, H M (1938) A simple method of estimating fifty per cent endpoints. Am J Hyg 27(3):493–497.

31. Nair A, Jacob S (2016) A simple practice guide for dose conversion between animals and human. J Basic Clin Pharm 7:27–31.

32. Ratanji KD, Derrick JP, Dearman RJ, Kimber I (2014) Immunogenicity of therapeutic proteins: Influence of aggregation. J Immunotoxicol 11(2):99–109.

33. Ngiow SF, von Scheidt B, Möller A, Smyth MJ, Teng MWL (2013) The interaction between murine melanoma and the immune system reveals that prolonged responses predispose for autoimmunity. Oncoimmunology 2(2):e23036.

34. Pardoll DM (2012) The blockade of immune checkpoints in cancer immunotherapy. Nat Rev Cancer 12(4):252–264.

35. Zou W, Wolchok JD, Chen L (2016) PD-L1 (B7-H1) and PD-1 pathway blockade for cancer therapy: Mechanisms, response biomarkers, and combinations. Sci Transl Med 8(328). doi:10.1126/scitranslmed.aad7118.

36. Brahmer JR, et al. (2010) Phase I study of single-agent anti-programmed death-1 (MDX-1106) in refractory solid tumors: Safety, clinical activity, pharmacodynamics, and immunologic correlates. J Clin Oncol 28(19):3167–3175.

37. Topalian SL, et al. (2012) Safety, activity, and immune correlates of anti–PD-1 antibody in cancer. N Engl J Med 366(26):9–19.

38. Padrón Á, et al. (2018) Age effects of distinct immune checkpoint blockade treatments in a mouse melanoma model. Exp Gerontol 105:146–154.

39. Hurez V, et al. (2012) Mitigating age-related immune dysfunction heightens the efficacy of tumor immunotherapy in aged mice. Cancer Res 72(8):2089–2099.

40. Nagai H, et al. (2004) In vivo elimination of CD25+ regulatory T cells leads to tumor rejection of B16F10 melanoma, when combined with interleukin-12 gene transfer. Exp Dermatol 13(10):613–620.

41. Jacobs JFM, Nierkens S, Figdor CG, Vries IJM De, Adema GJ (2012) Regulatory T cells in melanoma : the final hurdle towards effective immunotherapy ? Lancet Oncol 13:32–42.

42. Telang S, et al. (2011) Phase II trial of the regulatory T cell-depleting agent, denileukin diftitox, in patients with unresectable stage IV melanoma. BMC Cancer 11(515).

43. Vargas FA, et al. (2017) Fc-optimized anti-CD25 depletes tumor-infiltrating regulatory T cells and synergizes with PD-1 blockade to eradicate established tumors. Immunity 46(4):577–586.

44. Jacobs JFM, et al. (2010) Dendritic cell vaccination in combination with anti-CD25 monoclonal antibody treatment, a phase I/II study in metastatic melanoma patients. Clin Cancer Res. doi:10.1158/1078-0432.CCR-10-1757.

45. Foss FM (2000) DAB389IL-2 (Denileukin Diftitox, ONTAK): A New Fusion Protein Technology. Clin Lymphoma 1:S27–S31.

46. AM Pappenheimer Jr., Uchida T, Harper A (1972) An immunological study of the diphtheria toxin molecule. Immunochemistry 9:891–906.

